# Direct expression of CPT1a enables a high throughput platform for the discovery of CPT1a modulators

**DOI:** 10.1101/2024.12.15.628575

**Authors:** Jason Chen, Tuyen Tran, Anthony Wong, Luofei Wang, Pranavi Annaluru, Vibha Sreekanth, Samika Murthy, Laasya Munjeti, Tanya Park, Utkarsh Bhat, Glynnis Leong, Yumeng Li, Simeng Chen, Natalie Kong, Rushika Raval, Yining Xie, Shreya Somani, Aditi Bhambhani, Zoey Zhu, Landen Chu, Kimai Dosch, Edward Njoo, Zhan Chen

## Abstract

Carnitine palmitoyltransferase 1 (CPT1), which catalyzes the rate-limiting step of fatty acid oxidation, has been implicated in therapeutic approaches to several human diseases characterized by aberrant lipid metabolism. Isoform-specific quantification of CPT1 activity is essential in the characterization of small molecule inhibitors of CPT1, but several existing means to quantify enzymatic activity, including the use of radioisotope labeled carnitine, are not amenable to scalable, high throughput screening. Here, we demonstrate that mitochondrial extracts from Expi293 cells transfected with a CPT1a plasmid are a reliable and robust source of catalytically active human CPT1. Moreover, with a source of catalytically active enzyme in hand, we modified a previously reported colorimetric method of coenzyme A (CoA) easily scalable to a 96-well format for the screening of CPT1a inhibitors. This assay platform was validated by two previously reported inhibitors of CPT1a: *R*-etomoxir and perhexiline. To further demonstrate the applicability of this method in small molecule screening, we prepared and screened a library of 87 known small molecule APIs, validating the inhibitory effect of chlorpromazine on CPT1.

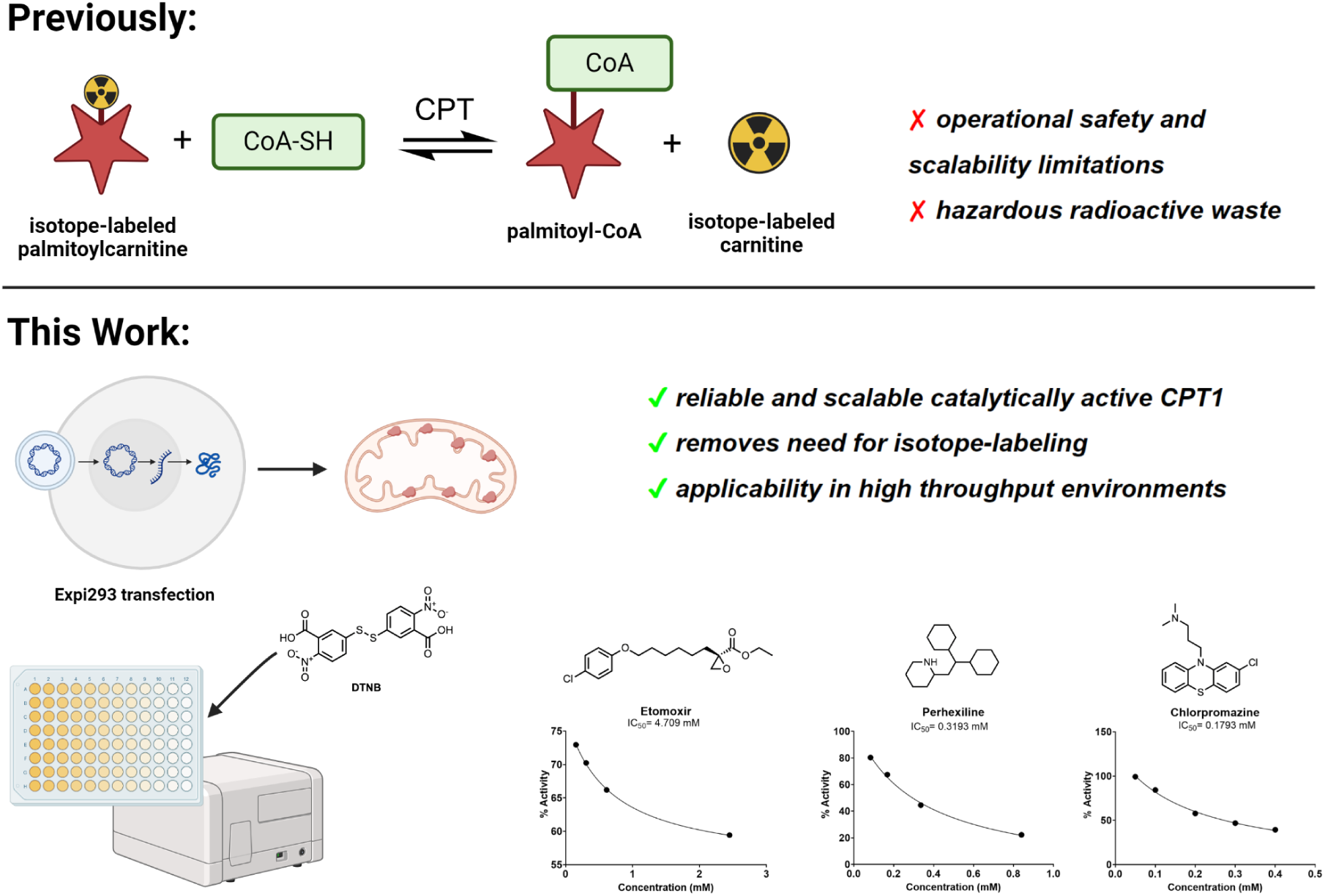

## Introduction

Carnitine palmitoyltransferase 1 (CPT1) catalyzes the rate-limiting step of fatty acid oxidation by shuttling long-chain fatty acids across the mitochondrial membrane [1,2]. This enzyme has been demonstrated to be a druggable target for the treatment of type 2 diabetes [3–5], cancer [6–9], obesity [10,11], and inflammation [12–14]. Carnitine acyltransferases catalyze the reversible transfer of acyl groups from acyl-coenzyme A (acyl-CoA) esters to L-carnitine, forming acylcarnitine esters [15–16]. The CPT system is made up of two separate proteins: CPT1, located in the outer mitochondrial membrane, is predominantly responsible for the forward acyl transfer to acylcarnitine, and CPT2, located in the inner mitochondrial membranes, catalyzes the hydrolysis of acylcarnitine back to acyl-CoA esters into the mitochondrial lumen [17–19].

To date, three distinct and yet closely related isoforms of CPT1 have been identified that are dissimilarly expressed in different tissues. CPT1a is known as the liver isoform, and is primarily expressed in brain, intestine, kidney, lung, ovary, pancreas, and spleen tissues [18,20]. CPT1a is endogenously inhibited by malonyl-CoA, the first intermediate in fatty acid synthesis [21,22]. CPT1b is predominantly found in high abundance in skeletal muscle, heart, and brown adipose tissue (BAT) [23]. Unlike CPT2, which has a well established structural basis from X-ray crystallography, no crystal structures of CPT1 exist to date, which further obfuscates efforts to understand the structural basis for small molecule ligands [24,25]. The third isoform, CPT1c, is predominantly expressed in the brain [26–28].

Given the clinical significance of fatty acid metabolism regulation in several indications [29–32], the CPT1 system has been subject to several small molecule inhibitor discovery campaigns [33]. Notably, etomoxir [34–36], a lipid-like α-epoxy carboxylate small molecule, has been previously identified as a covalent inhibitor of the active site of CPT1 for the treatment of type 2 diabetes mellitus [37] and chronic heart failure [38], but was withdrawn from late stage clinical studies due to severe off-target toxicity [39]. Based on such a background, a more selective CPT1 inhibitor named ST1326 (Teglicar) [40,41] was developed as a competitive, reversible, and isoform-selective CPT1 inhibitor with an improved toxicity and PK profile. Teglicar was in phase 2 studies [42] for the treatment of type 2 diabetes and has also shown great potential in the treatment of leukemias [43,44]. Other small molecule inhibitors, including perhexiline [45–48], dexamethasone [49,50], amiodarone [42, 51, 52], metoprolol [53,54], trimetazidine [55–57], and oxfenicine [58–60] have also been described to modulate activity of the CPT1 system.

The success of such small molecule campaigns in identifying lead compounds requires a sensitive and specific CPT1 activity assay. Traditionally, the catalytic activity of CPT1 has been measured through tritium-labeled L-[^3^H]carnitine [61–66]. While exceptionally sensitive, such tritium-based biochemical assays have inherent limitations in operational safety and scalability given the hazards of tritium waste and limited access to the equipment necessary to measure radiolabeled samples. Additionally, while several immunohistochemical approaches to quantify CPT1a concentration exist, specific immunohistochemical assays to measure the enzyme’s activity are not commercially available. Moreover, given the necessity of being in a mitochondrial-membrane environment for proper CPT1a structure and function, recombinant, commercially available sources of CPT1a are typically catalytically inactive, rendering them useless for inhibitor screening.

Rapid and sensitive spectrophotometric and fluorogenic assays for detection of carnitine palmitoyltransferase activity by measurement of released CoA-SH have been previously described [67–70]. Bieber et al. reported that the amount of reduced CoA liberated from palmitoyl-CoA by CPT is quantitated using 5,5′-dithiobis-(2-nitrobenzoic acid) (DTNB). This assay measures the initial rates of CoA-SH formation by CPT1-catalyzed deacylation of palmitoyl-CoA. Importantly, Bieber et al. demonstrate that this assay method does not require purification of recombinant human CPT1, but rather crude extracts from Sprague-Dawley rat liver could be directly employed as a reliable and standardizable source of human CPT1 [67,71]. While this approach circumvented the isolation instability of CPT1 as is typical of membrane-bound proteins, the *in vivo* source of mitochondrial extracts described therein are not easily sourceable and also potentially contain mixtures of CPT isoforms, among other enzymes present in fresh liver tissue [72].

To address the limited availability of high throughput screening platforms for evaluation of CPT1a activity and its inhibitors, we directly expressed CPT1a as a single isoform through transformation of Expi293 cells and show that the mitochondrial extracts isolated from *in-vitro* cell culture are a scalable and reliable source of catalytically active CPT1a. Furthermore, we modified a direct, colorimetric CoA detection method for CPT enzyme activity inspired by previous reports from Bieber et al. employing thiol-disulfide exchange from free CoA and DTNB, which leads to the production of 2-thio-5-nitrobenzoic acid (TNB) and an optical readout at 412 nm. We applied this system to the detection of free thiols from the release of CoA from palmitoyl-CoA. This assay detects the concentration of CoA liberated from CPT1 catalysis without the need to purify recombinant human CPT1 and was validated with a series of previously established CPT1 inhibitors. Further, to demonstrate the generalizability of our approach in high throughput small molecule screening campaigns, we applied this platform to screen for potential inhibitors of CPT1 in a sample of 87 small molecules. In accordance with previous reports, chlorpromazine was identified as a part of this high throughput screen to exert inhibitory effects on CPT1 [52, 73, 74]. The inhibitory effects we viewed in our high throughput screen demonstrate the validity of the modified assay, even when performed in large quantities.

## Materials and Methods

### Cell Culture

Mammalian Expi293F cells (Gibco, Cat. #A14527) were stably transfected with a human CPT1a expression plasmid with an ExpiFectamine 293 Kit (Gibco, Cat. #A14524) and maintained in Expi293 Expression Medium (Gibco, Cat. #12338018). All cells were cultured in 250 mL bottles, in a humidified incubator at 37°C in 5% CO_2_.

### Human CPT1a enzyme expression

To construct a human CPT1a expression plasmid, two expression constructions, CPT1A(NM_001876.3) ORF, which contains the whole human CPT1a sequence, and sp_CPT1A(NM_001876.3) ORF, which contains an N-terminal signal peptide for extracellular release, were each cloned into the vector pcDNA3.1+/C-(K)DYK backbone and purchased from GenScript USA Inc. The plasmid was transformed into TOP10 bacteria and amplified. After plasmid extraction and purification, the inserted CPT1a gene was sequenced. After sequencing confirmation, they were transfected into mammalian Expi293 cell lines. After growing the Expi293 culture, the supernatant and cell pellet were lysed with radioimmunoprecipitation assay (RIPA) buffer, and the protein was harvested. Protein concentration was quantified with a Bradford protein assay kit.

### Immunoassay to quantify CPT1a

ELISA was used to detect the expression level of human CPT1a in the supernatant and pellet, and validate successful transgenic expression of human CPT1a. Briefly, the CPT1a transfected cell pellet and supernatant, which contained 100 μg of total protein, was coated in a coating buffer (NaHCO_3_-Na_2_CO_3_ buffer, pH = 6.4) that was freshly made and aliquoted into 100 μL per well, and was left at 4°C for 18 hours. Following three washes with PBST (Phosphate buffered saline with Tween), 200 μL of blocking buffer was added to each well (2.5% w/v non-fat dry milk powder in PBST, 0.1% Tween-20, VWR Life Science, Cat. #0777-1L) for 1 hour at 18 °C. Following aspiration, mouse anti-human CPT1a monoclonal antibody (Abcam, Cat. #8F6AE9) was added in a 1:1,000 dilution (100 μL per well) and shaken for 2 hours at 18°C. Following a second wash, the goat-anti-mouse HRP secondary antibody (Sino Biological, Cat. #SSA007) was added with the blocking buffer. Subsequently, we added 100 μL of the diluted secondary antibody to each well after washing three times with PBST for 1 hour at 18°C while shaking. Then, 50 μL TMB (3,3’,5,5’-Tetramethylbenzidine) substrate (0.4 g/L, Tribioscience, Cat. #TBS5021) was added to each well after washing three times with PBST at 18°C. 100 μL of stop solution (1 M H_2_SO_4_) was then added to each well. We measured absorbance at 450 nm after 10 seconds of shaking. The Expi293 cell pellet protein was used as an endogenous control.

### CPT enzyme activity assay and inhibitor sensitivity assay

CPT spectrophotometric assay using DTNB was modified from the methods originally reported by Bieber et al. The reaction master mix was prepared with 35 μM Palmitoyl-CoA (Sigma Aldrich, Cat. #P9716), 1.5 mM EDTA (Acros, Cat. #AC118432500), 40 mM HEPES (ThermoScientific, Cat. #15630080), 1.25 mM L-carnitine (AK Scientific, Cat. #J94077), and a stock aliquot of the CPT1-containing mitochondrial extract. 100 mM Tris buffer (pH = 7.5) was used to make the final volume 90 μL. Samples were incubated at 30°C on a Compact Digital Dry Bath/Block Heater (ThermoFisher) for 15 min, at which point, 10 μL of 1 mg/mL DTNB (AK Scientific, Cat. #J92390) in 100 mM Tris buffer was added to each well. The plate was mixed on a plate shaker for 10 minutes and absorbances were recorded at 412 nm (Molecular Devices SpectraMAX). To correlate the appearance of yellow with the absolute concentration of CoA and to determine the dynamic range of the assay, we prepared a standard curve.

Two previously reported CPT1 inhibitors, etomoxir sodium salt (Cayman Chemical, Cat. #11969) and perhexiline (AK Scientific, Cat. #X4611), were used to validate the potential of this assay as a platform to screen for novel CPT1 inhibitors. IC_50_ calculations were performed on the GraphPad Prism 10 software.

### High-throughput screening of a small molecule library

To demonstrate the validity of this approach in scalable, high-throughput screening campaigns, 87 small molecule compounds were screened using this platform, with etomoxir as a positive control. The compounds screened were selected from a library of natural products, anti-virals, immunomodulators, and other bioactive compounds. Compounds were diluted to 10 mM in dimethyl sulfoxide (DMSO) (Stellar Chemical, 99.99%) and tested at a final concentration of 0.556 mM. Compounds that exhibited inhibitory effects were run in a serial concentration to create an inhibitory effect curve and determine the IC_50_.

### Data analysis

All data and statistics (mean, SEM, p-values) were analyzed with GraphPad Prism 10 and Excel. Acquired data are presented as mean ± SEM. P-values of 0.05 were considered statistically significant.

## Results

### Human CPT1a enzyme expression and protein isolation

Two human CPT1a gene expression constructions, pcDNA3.1+/C-(K)DYK with CPT1A(NM_001876.3) ORF Clone and sp_CPT1A(NM_001876.3) ORF Clone, were sequenced and aligned to Genbank. The sequences are identical to previously deposited sequences on Genbank. The supernatant and cell pellet proteins were harvested separately after transfection into mammalian cell Expi293 as shown in **Figure 1**. The normal Expi293 cell pellet proteins were harvested as a control.

**Figure 1.**
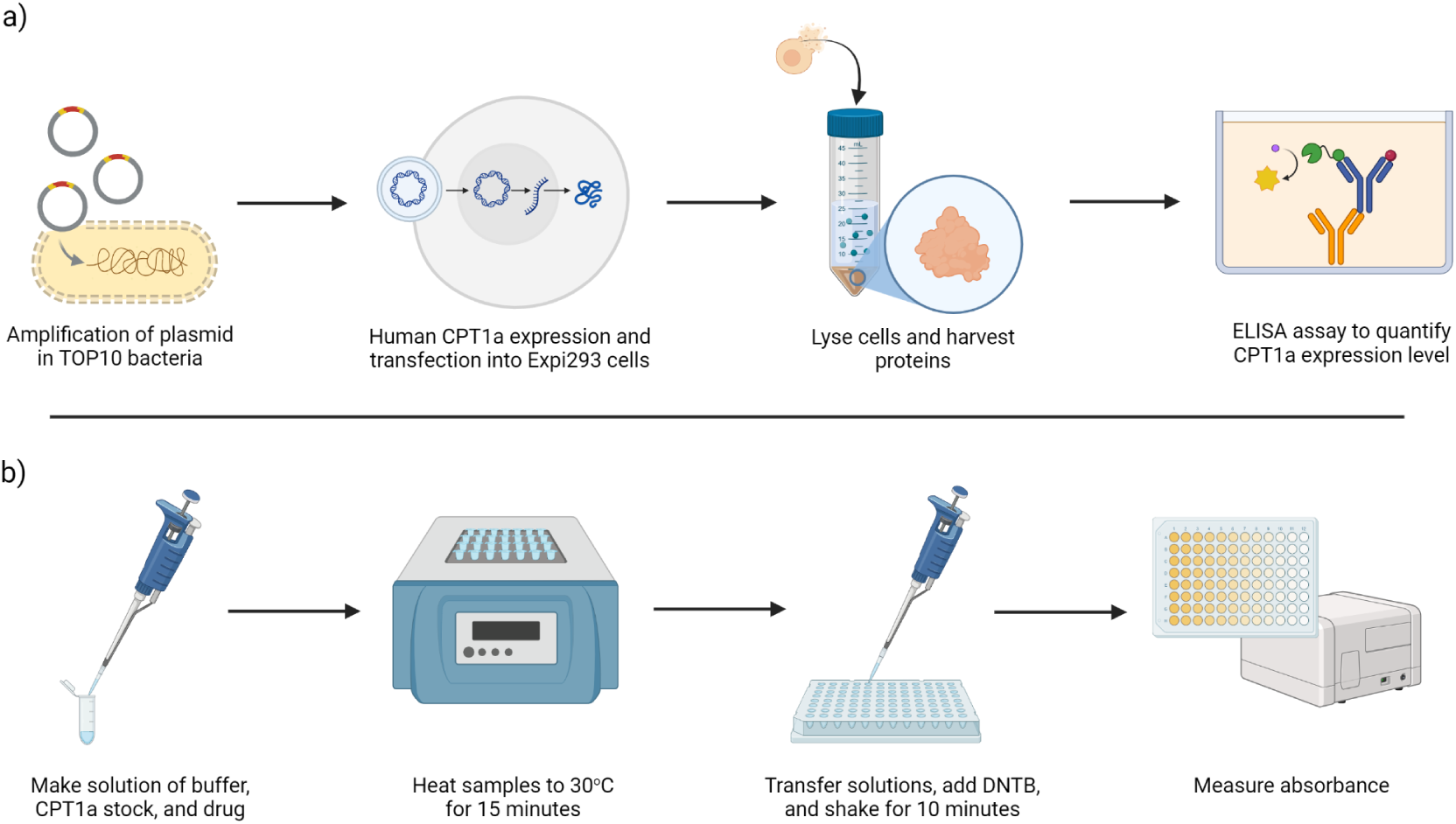
Workflow of CPT1 enzyme expression, protein purification, and activity assay using DTNB. a) The workflow of purification and quantification of human CPT1a enzyme. First, the plasmid was constructed and transformed into TOP10 bacteria for amplification. Plasmids were extracted and purified before transfected into Expi293 cells, where they were cultured. Cells were then lysed, and the supernatant and pellet were collected for protein quantification by ELISA. b) The CPT1 enzyme activity assay workflow. First, a solution of the buffer reagents, diluted CPT1 stock, and drug was created. The samples were mixed well and warmed to 30°C with a block plate heater for 15 minutes for the reaction to occur. Samples were then transferred to 96-well plates and DTNB was added. The plate was shaken on a cell shaker for 10 minutes before the absorbance was read in a plate reader at 412 nm.

### CPT1a immunogenicity assay

ELISA was used to check the immunity and protein expression level of CPT1a in transfected Expi293 mitochondria. The CPT1 expression level in the CPT1a transgenic cell pellet had a 13-fold increase of control cell pellet, while minimal CPT1 was detected in the supernatant, as shown in **Figure 2**. The supernatant concentration of CPT1a in cells transfected with a signal peptide fusion protein did not exhibit higher extracellular content of CPT1a, suggesting that CPT1a cannot be secreted out of the cell into the supernatant, potentially because it is very tightly bound to the mitochondrial membrane.

**Figure 2.**
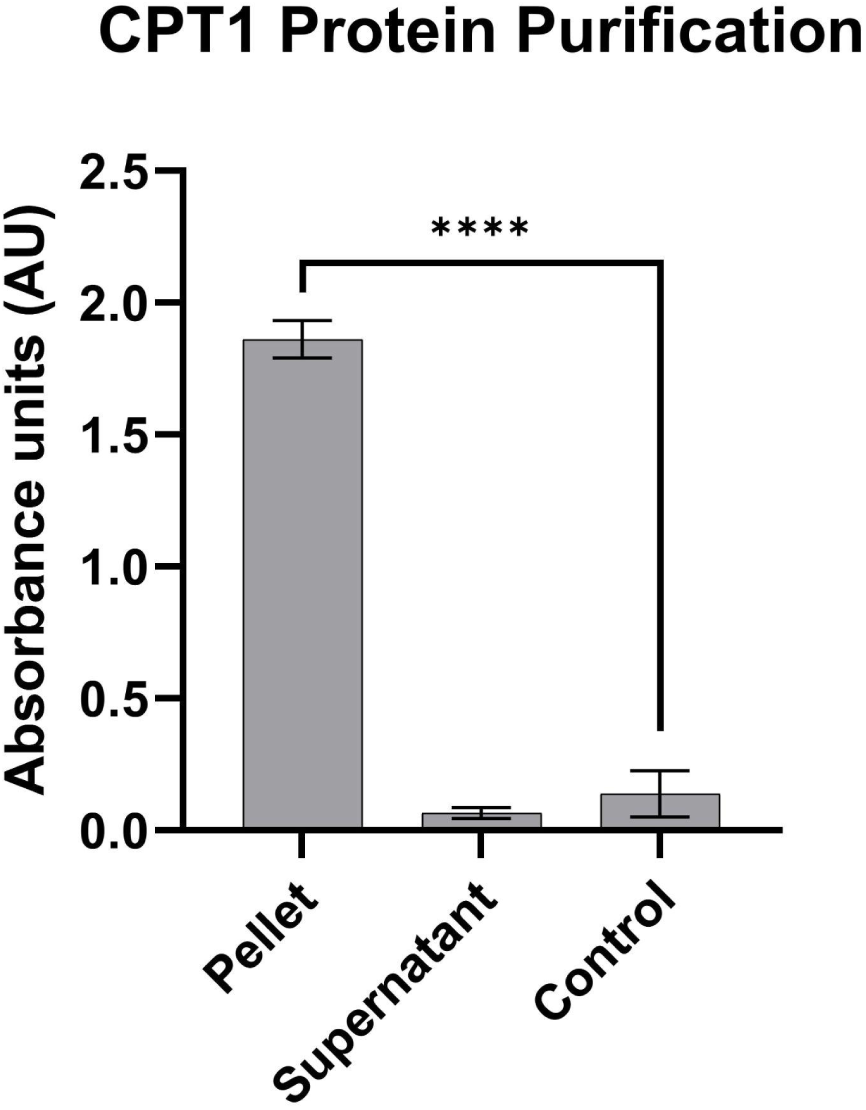
ELISA of CPT1a transgenic cell line. CPT1a immunogenicity assay using ELISA. Pellet is the human CPT1a transgenic cell line pellet. Supernatant is the human CPT1a transgenic cell line supernatant. Control is the wildtype Expi293 cell pellet without CPT1a gene transfection. The data is represented as mean ± SEM (n = 4), and significance was calculated using a Welch’s t-test (**P < 0.01, ****P<0.0001).

### CPT1 enzyme activity assay validation

A CoA standard curve with authentic CoA (Cayman Chemical, Cat. #16147) was used to quantify the concentration of CoA being released during the catalysis. The enzyme activity calculation formula: 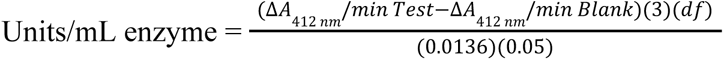, where: 3 =Total volume (in milliliters) of assay, df = Dilution Factor, 0.0136 = Micromolar extinction coefficient for TNB at 412 nm, 0.05 = Volume (in milliliters) of enzyme used. As a background, the CPT1a cell pellet has a value of 5.02 ± 0.2 μM.

The value of CPT1 enzyme activity is proportional to the concentration of CPT1 enzyme as shown in **Figure 3**. The inhibition efficacy of etomoxir was calculated by running a serial dilution of etomoxir concentrations in the enzyme activity assay. Replicates of this assay demonstrated nearly identical etomoxir-dependent activity profiles, indicating a high degree of reliability.

**Figure 3.**
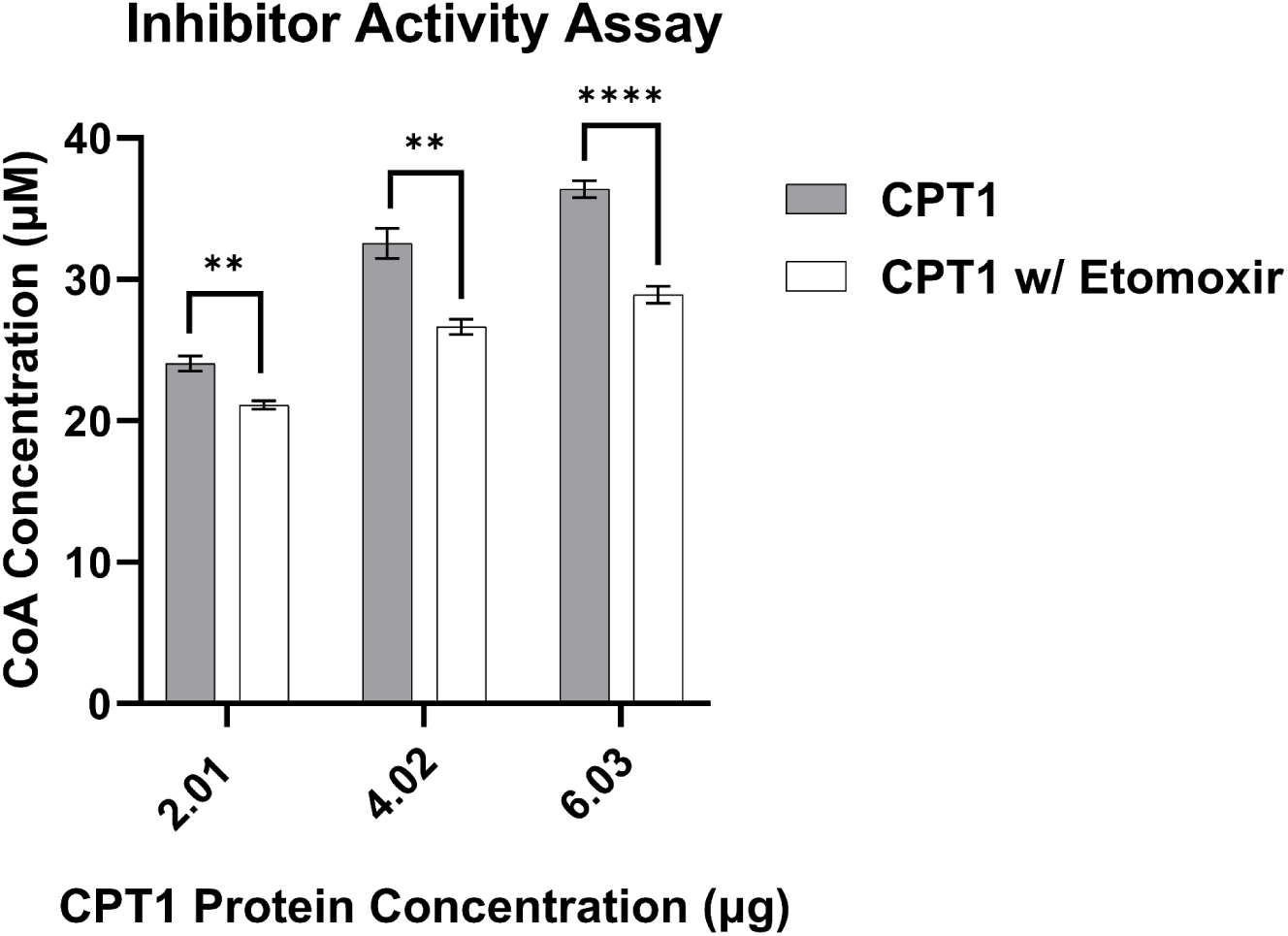
CPT activity assay and inhibitor sensitivity assay with varying enzyme concentrations. CPT activity assay and etomoxir inhibitor sensitivity assay. We used 3 μL, 6 μL, and 9 μL of the same cell lysis pellet solution with a protein concentration of 0.67 μg/μL. 2 μL of a 15.58 mM stock solution of etomoxir was added to each group to make a final concentration of 0.306 mM. Data is represented as mean ± SEM (n = 3), and significance was calculated using a Welch’s t-test (**P < 0.01, ****P<0.0001).

### CPT1 enzyme inhibitor sensitivity assay

We used the CPT1 enzyme inhibition assay to validate the new protocols that were shown in **Figure 3** and **Figure 4**. The inhibition rate IC_50_ of etomoxir was 4.709 mM (**Figure 5**). Another inhibitor, perhexiline, was additionally used to validate this platform. Both inhibition curves of etomoxir and perhexiline indicate that the new CPT1 enzyme activity assay method can quantify the different effects among different inhibitors.

**Figure 4.**
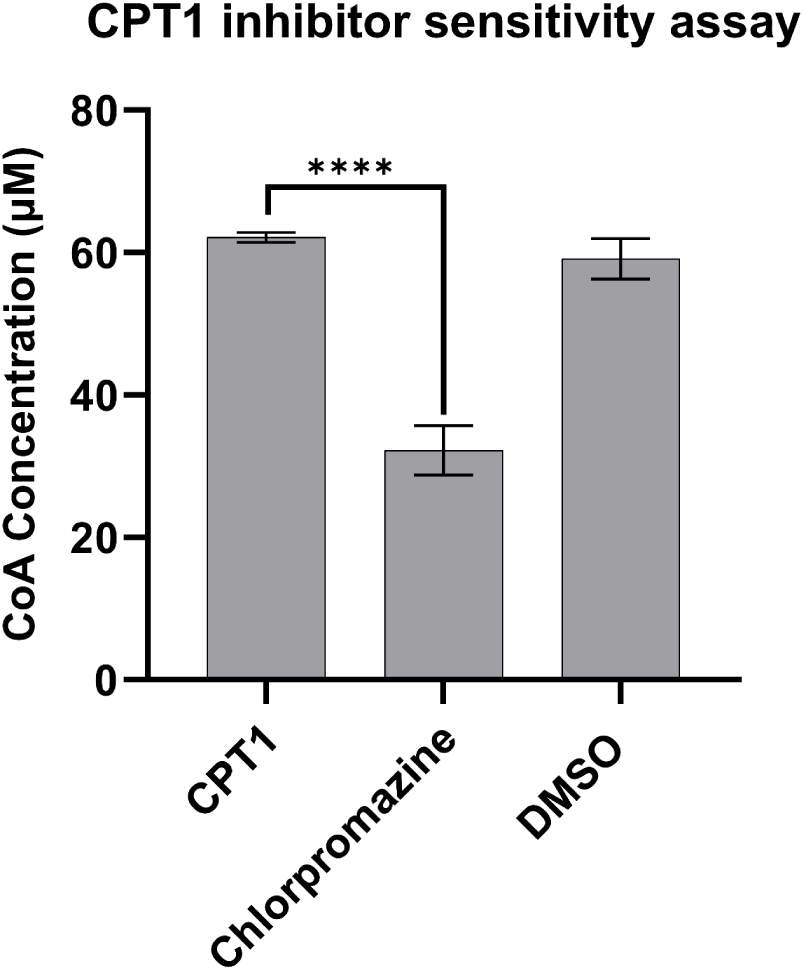
CPT1 inhibitor sensitivity assay. CPT1 inhibitor sensitivity assay using chlorpromazine. We used the same amount of human CPT1a pellet solution at the protein concentration of 0.67 μg/μL, 2 μL each group. 2 μL of 10 mM chlorpromazine as a solution in DMSO was used. 2 μL of DMSO (Fisher Scientific, ≥ 99.7%) was used as a negative control. Data is represented as means ± SEM (n = 3), and significance was calculated using a Welch’s t-test (**P < 0.01, ****P < 0.0001).

**Figure 5.**
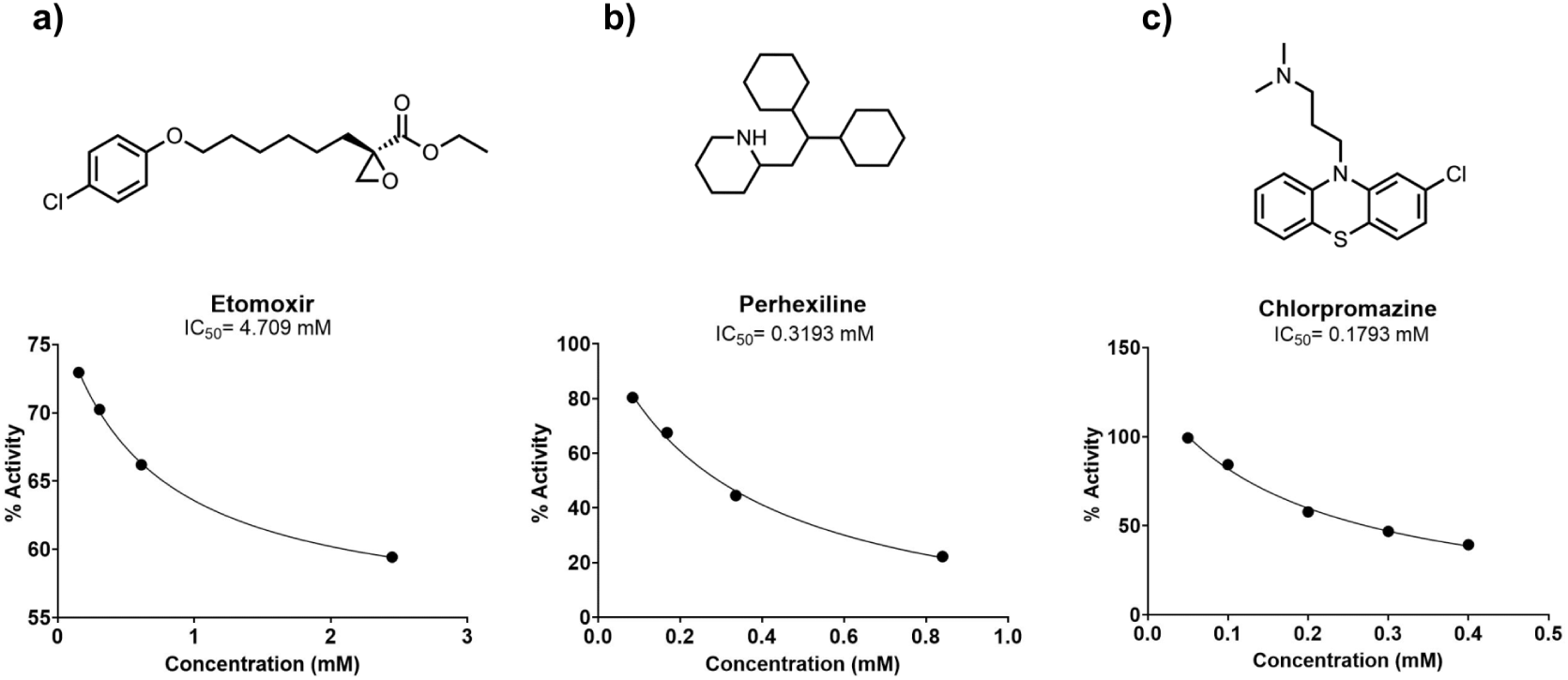
IC_50_ of etomoxir, perhexiline, and chlorpromazine. IC_50_ of a) etomoxir, b) perhexiline, c) chlorpromazine. A series of concentrations of each compound was used accordingly. We used the same amount of human CPT1a pellet solution with a protein concentration of 0.67 μg/μL, adding 2 μL of protein in each group.

### Application and performance qualification to screen small molecule compounds

In order to demonstrate the applicability of this system in the high-throughput screening of new CPT1 inhibitors and activators, we prepared and screened a library of 87 small molecule compounds using the new approaches, including several FDA-approved small molecules for different indications, as well as natural products and previously reported preclinical candidates. From this screen, we validated that chlorpromazine, a previously clinically studied antipsychotic small molecule, exhibits inhibitory activity against CPT1a, as shown in **Figures 4-5**. Its IC_50_ for CPT1a inhibition is 0.1793 mM.

## Discussion

CPT1 facilitates the transfer of long chain fatty acids into the mitochondria for beta-oxidation, and a lack of regulation promotes the development of various human diseases. Therefore, a highly reproducible, cost-effective method of identifying CPT1 inhibitors paves the way for substantial medical breakthroughs in developing potential treatments for type 2 diabetes and cancer. While highly sensitive and accurate radioisotope-based quantification methods exist for the direct measurement of CPT1a activity, safety limitations and limited access to scintillation equipment pose major drawbacks to the applicability and scalability of ^3^H-carnitine based assays. Additionally, while commercially available ELISA kits exist for detection of CPT1 through immunogenicity with poly- or monoclonal antibodies, such methods cannot detect the enzyme activity of CPT1.

In summary, we modified and optimized a highly robust assay for the quantification of CPT1 activity. This method allows for the use of the direct cell lysis of human CPT1 transformed Expi293 cells without the need for purification of recombinant proteins. Further, we optimized the spectrophotometric detection of CoA liberated from CPT1 catalysis using a well-established DTNB-based detection of free thiols, which is a proxy for enzyme activity based on the direct relationship between CPT1 catalytic activity and increasing CoA concentration. We then validated this approach by using previously reported CPT1 inhibitors, etomoxir, perhexiline, and chlorpromazine, and found that this assay method provided highly reproducible and dose-responsive quantification for CPT1 activity.

Our approach overcomes the limitations of isotope-based screening methods and is thus suitable for high-throughput applications. Finally, we show that this assay is easily adaptable to a 96-well format, suggesting the feasibility of performing screens on large sets of samples, including the potential for automation. The IC_50_ of chlorpromazine we reported using this assay is consistent with the IC_50_ trends published in the first report of chlorpromazine as a CPT1 inhibitor, which used the isotope forward exchange method, further demonstrating the validity of this assay even in high throughput formats [73].

Since the reaction of DTNB with the sulfhydryl group on CoA results in an increase in A_412nm_ due to the yellow TNB anion, compounds that absorb in a similar range will interfere with the assay and the data acquisition, and thus are not suitable for screening by this method. This strategy has potential to be further applied to the study of small molecule ligands to other CPT1 isoforms.

## CRediT authorship contribution statement

**Jason Chen:** Data curation, Investigation. **Tuyen Tran:** Investigation, Resources. **Anthony Wong:** Data curation, Investigation, Validation, Writing – review & editing. **Luofei Wang:** Data curation, Investigation, Validation, Writing – review & editing. **Pranavi Annaluru:** Data curation, Investigation, Methodology, Validation, Writing – review & editing. **Vibha Sreekanth:** Data curation, Investigation, Methodology, Validation, Writing – review & editing. **Samika Murthy:** Data curation, Investigation, Methodology, Validation, Writing – review & editing. **Laasya Munjeti:** Data curation, Investigation, Validation, Writing – review & editing. **Tanya Park:** Data curation, Investigation, Validation, Writing – review & editing. **Utkarsh Bhat:** Data curation, Investigation, Methodology, Validation, Writing – review & editing. **Glynnis Leong:** Data curation, Investigation, Validation, Writing – review & editing. **Yumeng Li:** Data curation, Investigation, Methodology, Validation, Writing – review & editing. **Simeng Chen:** Data curation, Investigation, Methodology. **Natalie Kong:** Formal analysis, Resources, Visualization, Writing – original draft, Writing – review & editing. **Rushika Raval:** Formal analysis, Resources, Visualization, Writing – original draft, Writing – review & editing. **Yining Xie:** Formal analysis, Resources, Visualization, Writing – original draft, Writing – review & editing. **Shreya Somani:** Resources. **Aditi Bhambhani:** Data curation, Investigation, Validation, Writing – review & editing. **Zoey Zhu:** Data curation, Investigation, Validation, Writing – review & editing. **Landen Chu**: Data curation, Investigation, Validation, Writing - Review & Editing. **Kimai Dosch:** Resources. **Edward Njoo:** Conceptualization, Funding acquisition, Project administration, Resources, Supervision, Writing – original draft, Writing – review & editing. **Zhan Chen:** Conceptualization, Funding acquisition, Project administration, Supervision, Writing – original draft, Writing – review & editing.

