## Supporting Information Document for "Direct expression of CPT1a enables a high throughput platform for the discovery of CPT1a modulators"

A. Non-fusion Plasmid SI-1
   a. Non-fusion Plasmid Sequence SI-1
   b. Non-fusion Plasmid Map SI-1
B. Fusion Plasmid Sequence and Map SI-2
   a. Fusion Plasmid Sequence SI-2
   b. Fusion Plasmid Map SI-2
C. Standard Curve for DTNB SI-3
D. 78 Compound Library SI-4

### A. Non-fusion Plasmid

#### a. Non-fusion Plasmid Sequence

pcDNA3.1

> CPT1A(NM_001876.3) ORF Clone

ATGGCAGAAGCTCACCAAGCTGTGGCCTTTCAGTTCACGGTCACTCCGGACGGGATTGACCT
GCGGCTGAGCCATGAAGCTCTTAGACAAATCTATCTCTCTGGACTTCATTCCTGGAAAAAGAA
GTTCATCAGATTCAAGAACGGCATCATCACTGGCGTGTACCCGGCAAGCCCCTCCAGTTGGCT
TATCGTGGTGGTGGGCGTGATGACAACGATGTACGCCAAGATCGACCCCTCGTTAGGAATAAT
TGCAAAAATCAATCGGACTCTGGAAACGGCCAACTGCATGTCCAGCCAGACGAAGAACGTGG
TCAGCGGCGTGCTGTTTGGCACCGGCCTGTGGGTGGCCCTCATCGTCACCATGCGCTACTCCC
TGAAAGTGCTGCTCTCCTACCACGGGTGGATGTTCACTGAGCACGGCAAGATGAGTCGTGCC
ACCAAGATCTGGATGGGTATGGTCAAGATCTTTTCAGGCCGAAAACCCATGTTGTACAGCTTC
CAGACATCGCTGCCTCGCCTGCCGGTCCCGGCTGTCAAAGACACTGTGAACAGGTATCTACA
GTCGGTGAGGCCTCTTATGAAGGAAGAAGACTTCAAACGGATGACAGCACTTGCTCAAGATT
TTGCTGTCGGTCTTGGACCAAGATTACAGTGGTATTTGAAGTTAAAATCCTGGTGGGCTACAA
ATTACGTGAGCGACTGGTGGGAGGAGTACATCTACCTCCGAGGACGAGGGCCGCTCATGGTG
AACAGCAACTATTATGCCATGGATCTGCTGTATATCCTTCCAACTCACATTCAGGCAGCAAGAG
CCGGCAACGCCATCCATGCCATCCTGCTTTACAGGCGCAAACTGGACCGGGAGGAAATCAAA
CCAATTCGTCTTTTGGGATCCACGATTCCACTCTGCTCCGCTCAGTGGGAGCGGATGTTTAATA
CTTCCCGGATCCCAGGAGAGGAGACAGACACCATCCAGCACATGAGAGACAGCAAGCACATC
GTCGTGTACCATCGAGGACGCTACTTCAAGGTCTGGCTCTACCATGATGGGCGGCTGCTGAAG
CCCCGGGAGATGGAGCAGCAGATGCAGAGGATCCTGGACAATACCTCGGAGCCTCAGCCCGG
GGAGGCCAGGCTGGCAGCCCTCACCGCAGGAGACAGAGTTCCCTGGGCCAGGTGTCGTCAG
GCCTATTTTGGACGTGGGAAAAATAAGCAGTCTCTTGATGCTGTGGAGAAAGCAGCGTTCTTC
GTGACGTTAGATGAAACTGAAGAAGGATACAGAAGTGAAGACCCGGATACGTCAATGGACAG
CTACGCCAAATCTCTACTACACGGCCGATGTTACGACAGGTGGTTTGACAAGTCGTTCACGTT
TGTTGTCTTCAAAAACGGGAAGATGGGCCTCAACGCTGAACACTCCTGGGCAGATGCGCCGA
TCGTGGCCCACCTTTGGGAGTACGTCATGTCCATTGACAGCCTCCAGCTGGGCTATGCGGAGG
ATGGGCACTGCAAAGGCGACATCAATCCGAACATTCCGTACCCCACCAGGCTGCAGTGGGAC
ATCCCGGGGGAATGTCAAGAGGTTATAGAGACCTCCCTGAACACCGCAAATCTTCTGGCAAA
CGACGTGGATTTCCATTCCTTCCCATTCGTAGCCTTTGGTAAAGGAATCATCAAGAAATGTCGC
ACGAGCCCAGACGCCTTTGTGCAGCTGGCCCTCCAGCTGGCGCACTACAAGGACATGGGCAA
GTTTTGCCTCACATACGAGGCCTCCATGACCCGGCTCTTCCGAGAGGGGAGGACGGAGACCG
TGCGCTCCTGCACCACTGAGTCATGCGACTTCGTGCGGGCCATGGTGGACCCGGCCCAGACG
GTGGAACAGAGGCTGAAGTTGTTCAAGTTGGCGTCTGAGAAGCATCAGCATATGTATCGCCTC
GCCATGACCGGCTCTGGGATCGATCGTCACCTCTTCTGCCTTTACGTGGTGTCTAAATATCTCG
CTGTGGAGTCCCCTTTCCTTAAGGAAGTTTTATCTGAGCCTTGGAGATTATCAACAAGCCAGA
CCCCTCAGCAGCAAGTGGAGCTGTTTGACTTGGAGAATAACCCAGAGTACGTGTCCAGCGGA
GGGGGCTTTGGACCGGTTGCTGATGACGGCTATGGTGTGTCGTACATCCTTGTGGGAGAGAAC
CTCATCAATTTCCACATTTCTTCCAAGTTCTCTTGCCCTGAGACGGATTCTCATCGCTTTGGAA
GGCACCTGAAAGAAGCAATGACTGACATCATCACTTTGTTTGGTCTCAGTTCTAATTCCAAAA
AG

#### b. Non-fusion Plasmid Map

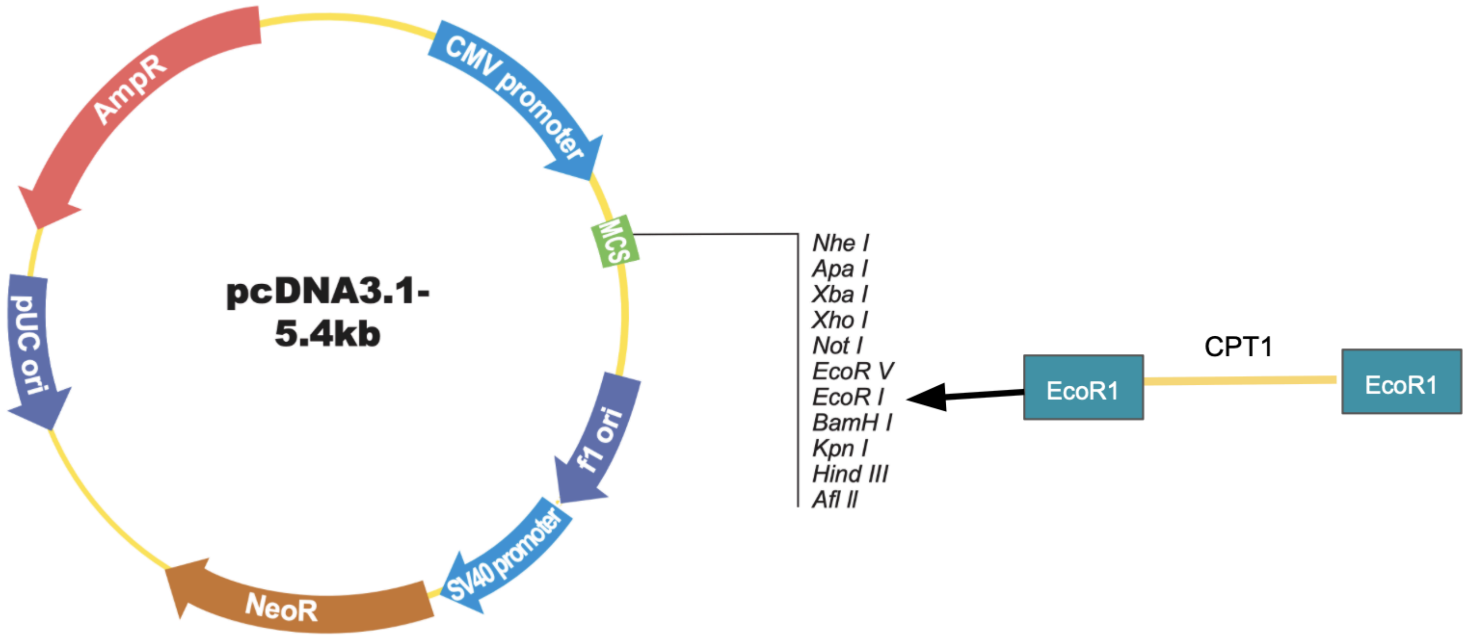

### B. Fusion Plasmid

#### a. Fusion Plasmid Sequence

>CPT1A(NM_001876.3) ORF Clone
GACGGATCGGGAGATCTCCCGATCCCCTATGGTGCACTCTCAGTACAATCTGCTCTGATGCCGC
ATAGTTAAGCCAGTATCTGCTCCCTGCTTGTGTGTTGGAGGTCGCTGAGTAGTGCGCGAGCAA
AATTTAAGCTACAACAAGGCAAGGCTTGACCGACAATTGCATGAAGAATCTGCTTAGGGTTAG
GCGTTTTGCGCTGCTTCGCGATGTACGGGCCAGATATACGCGTTGACATTGATTATTGACTAGT
TATTAATAGTAATCAATTACGGGGTCATTAGTTCATAGCCCATATATGGAGTTCCGCGTTACATA
ACTTACGGTAAATGGCCCGCCTGGCTGACCGCCCAACGACCCCCGCCCATTGACGTCAATAAT
GACGTATGTTCCCATAGTAACGCCAATAGGGACTTTCCATTGACGTCAATGGGTGGAGTATTTA
CGGTAAACTGCCCACTTGGCAGTACATCAAGTGTATCATATGCCAAGTACGCCCCCTATTGACG
TCAATGACGGTAAATGGCCCGCCTGGCATTATGCCCAGTACATGACCTTATGGGACTTTCCTAC
TTGGCAGTACATCTACGTATTAGTCATCGCTATTACCATGGTGATGCGGTTTTGGCAGTACATCA
ATGGGCGTGGATAGCGGTTTGACTCACGGGGATTTCCAAGTCTCCACCCCATTGACGTCAATG
GGAGTTTGTTTTGGCACCAAAATCAACGGGACTTTCCAAAATGTCGTAACAACTCCGCCCCAT
TGACGCAAATGGGCGGTAGGCGTGTACGGTGGGAGGTCTATATAAGCAGAGCTCTCTGGCTAA
CTAGAGAACCCACTGCTTACTGGCTTATCGAAATTAATACGACTCACTATAGGGAGACCCAAG
CTGGCTAGCGTTTAAACTTAAGCTTGGTACCGAGCTCGGATCCGCCACCATGGGCTGGAGCTG
CATCATCCTGTTCCTGGTGGCCACAGCCACCGGCGTGCACTCTGCAGAAGCTCACCAAGCTGT
GGCCTTTCAGTTCACGGTCACTCCGGACGGGATTGACCTGCGGCTGAGCCATGAAGCTCTTAG
ACAAATCTATCTCTCTGGACTTCATTCCTGGAAAAAGAAGTTCATCAGATTCAAGAACGGCAT
CATCACTGGCGTGTACCCGGCAAGCCCCTCCAGTTGGCTTATCGTGGTGGTGGGCGTGATGAC
AACGATGTACGCCAAGATCGACCCCTCGTTAGGAATAATTGCAAAAATCAATCGGACTCTGGA
AACGGCCAACTGCATGTCCAGCCAGACGAAGAACGTGGTCAGCGGCGTGCTGTTTGGCACCG
GCCTGTGGGTGGCCCTCATCGTCACCATGCGCTACTCCCTGAAAGTGCTGCTCTCCTACCACG
GGTGGATGTTCACTGAGCACGGCAAGATGAGTCGTGCCACCAAGATCTGGATGGGTATGGTC
AAGATCTTTTCAGGCCGAAAACCCATGTTGTACAGCTTCCAGACATCGCTGCCTCGCCTGCCG
GTCCCGGCTGTCAAAGACACTGTGAACAGGTATCTACAGTCGGTGAGGCCTCTTATGAAGGA
AGAAGACTTCAAACGGATGACAGCACTTGCTCAAGATTTTGCTGTCGGTCTTGGACCAAGATT
ACAGTGGTATTTGAAGTTAAAATCCTGGTGGGCTACAAATTACGTGAGCGACTGGTGGGAGG
AGTACATCTACCTCCGAGGACGAGGGCCGCTCATGGTGAACAGCAACTATTATGCCATGGATC
TGCTGTATATCCTTCCAACTCACATTCAGGCAGCAAGAGCCGGCAACGCCATCCATGCCATCCT
GCTTTACAGGCGCAAACTGGACCGGGAGGAAATCAAACCAATTCGTCTTTTGGGATCCACGA
TTCCACTCTGCTCCGCTCAGTGGGAGCGGATGTTTAATACTTCCCGGATCCCAGGAGAGGAGA
CAGACACCATCCAGCACATGAGAGACAGCAAGCACATCGTCGTGTACCATCGAGGACGCTAC
TTCAAGGTCTGGCTCTACCATGATGGGCGGCTGCTGAAGCCCCGGGAGATGGAGCAGCAGAT
GCAGAGGATCCTGGACAATACCTCGGAGCCTCAGCCCGGGGAGGCCAGGCTGGCAGCCCTCA
CCGCAGGAGACAGAGTTCCCTGGGCCAGGTGTCGTCAGGCCTATTTTGGACGTGGGAAAAAT
AAGCAGTCTCTTGATGCTGTGGAGAAAGCAGCGTTCTTCGTGACGTTAGATGAAACTGAAGA
AGGATACAGAAGTGAAGACCCGGATACGTCAATGGACAGCTACGCCAAATCTCTACTACACG
GCCGATGTTACGACAGGTGGTTTGACAAGTCGTTCACGTTTGTTGTCTTCAAAAACGGGAAG
ATGGGCCTCAACGCTGAACACTCCTGGGCAGATGCGCCGATCGTGGCCCACCTTTGGGAGTA

CGTCATGTCCATTGACAGCCTCCAGCTGGGCTATGCGGAGGATGGGCACTGCAAAGGCGACA
TCAATCCGAACATTCCGTACCCCACCAGGCTGCAGTGGGACATCCCGGGGGAATGTCAAGAG
GTTATAGAGACCTCCCTGAACACCGCAAATCTTCTGGCAAACGACGTGGATTTCCATTCCTTC
CCATTCGTAGCCTTTGGTAAAGGAATCATCAAGAAATGTCGCACGAGCCCAGACGCCTTTGTG
CAGCTGGCCCTCCAGCTGGCGCACTACAAGGACATGGGCAAGTTTTGCCTCACATACGAGGC
CTCCATGACCCGGCTCTTCCGAGAGGGGAGGACGGAGACCGTGCGCTCCTGCACCACTGAGT
CATGCGACTTCGTGCGGGCCATGGTGGACCCGGCCCAGACGGTGGAACAGAGGCTGAAGTTG
TTCAAGTTGGCGTCTGAGAAGCATCAGCATATGTATCGCCTCGCCATGACCGGCTCTGGGATC
GATCGTCACCTCTTCTGCCTTTACGTGGTGTCTAAATATCTCGCTGTGGAGTCCCCTTTCCTTAA
GGAAGTTTTATCTGAGCCTTGGAGATTATCAACAAGCCAGACCCCTCAGCAGCAAGTGGAGC
TGTTTGACTTGGAGAATAACCCAGAGTACGTGTCCAGCGGAGGGGGCTTTGGACCGGTTGCT
GATGACGGCTATGGTGTGTCGTACATCCTTGTGGGAGAGAACCTCATCAATTTCCACATTTCTT
CCAAGTTCTCTTGCCCTGAGACGGATTCTCATCGCTTTGGAAGGCACCTGAAAGAAGCAATG
ACTGACATCATCACTTTGTTTGGTCTCAGTTCTAATTCCAAAAAGGATTACAAGGATGACGAC
GATAAGTGATAAACCCGCTGATCAGCCTCGACTGTGCCTTCTAGTTGCCAGCCATCTGTTGTTT
GCCCCTCCCCCGTGCCTTCCTTGACCCTGGAAGGTGCCACTCCCACTGTCCTTTCCTAATAAA
ATGAGGAAATTGCATCGCATTGTCTGAGTAGGTGTCATTCTATTCTGGGGGGTGGGGTGGGGC
AGGACAGCAAGGGGGAGGATTGGGAAGACAATAGCAGGCATGCTGGGGATGCGGTGGGCTC
TATGGCTTCTGAGGCGGAAAGAACCAGCTGGGGCTCTAGGGGGTATCCCCACGCGCCCTGTA
GCGGCGCATTAAGCGCGGCGGGTGTGGTGGTTACGCGCAGCGTGACCGCTACACTTGCCAGC
GCCCTAGCGCCCGCTCCTTTCGCTTTCTTCCCTTCCTTTCTCGCCACGTTCGCCGGCTTTCCCC
GTCAAGCTCTAAATCGGGGGCTCCCTTTAGGGTTCCGATTTAGTGCTTTACGGCACCTCGACC
CCAAAAAACTTGATTAGGGTGATGGTTCACGTAGTGGGCCATCGCCCTGATAGACGGTTTTTC
GCCCTTTGACGTTGGAGTCCACGTTCTTTAATAGTGGACTCTTGTTCCAAACTGGAACAACAC
TCAACCCTATCTCGGTCTATTCTTTTGATTTATAAGGGATTTTGCCGATTTCGGCCTATTGGTTAA
AAAATGAGCTGATTTAACAAAAATTTAACGCGAATTAATTCTGTGGAATGTGTGTCAGTTAGG
GTGTGGAAAGTCCCCAGGCTCCCCAGCAGGCAGAAGTATGCAAAGCATGCATCTCAATTAGT
CAGCAACCAGGTGTGGAAAGTCCCCAGGCTCCCCAGCAGGCAGAAGTATGCAAAGCATGCAT
CTCAATTAGTCAGCAACCATAGTCCCGCCCCTAACTCCGCCCATCCCGCCCCTAACTCCGCCCA
GTTCCGCCCATTCTCCGCCCCATGGCTGACTAATTTTTTTTATTTATGCAGAGGCCGAGGCCGC
CTCTGCCTCTGAGCTATTCCAGAAGTAGTGAGGAGGCTTTTTTGGAGGCCTAGGCTTTTGCAA
AAAGCTCCCGGGAGCTTGTATATCCATTTTCGGATCTGATCAAGAGACAGGATGAGGATCGTT
TCGCATGATTGAACAAGATGGATTGCACGCAGGTTCTCCGGCCGCTTGGGTGGAGAGGCTATT
CGGCTATGACTGGGCACAACAGACAATCGGCTGCTCTGATGCCGCCGTGTTCCGGCTGTCAGC
GCAGGGGCGCCCGGTTCTTTTTGTCAAGACCGACCTGTCCGGTGCCCTGAATGAACTGCAGG
ACGAGGCAGCGCGGCTATCGTGGCTGGCCACGACGGGCGTTCCTTGCGCAGCTGTGCTCGAC
GTTGTCACTGAAGCGGGAAGGGACTGGCTGCTATTGGGCGAAGTGCCGGGGCAGGATCTCCT
GTCATCTCACCTTGCTCCTGCCGAGAAAGTATCCATCATGGCTGATGCAATGCGGCGGCTGCAT
ACGCTTGATCCGGCTACCTGCCCATTCGACCACCAAGCGAAACATCGCATCGAGCGAGCACG
TACTCGGATGGAAGCCGGTCTTGTCGATCAGGATGATCTGGACGAAGAGCATCAGGGGCTCG
CGCCAGCCGAACTGTTCGCCAGGCTCAAGGCGCGCATGCCCGACGGCGAGGATCTCGTCGTG
ACCCATGGCGATGCCTGCTTGCCGAATATCATGGTGGAAAATGGCCGCTTTTCTGGATTCATCG
ACTGTGGCCGGCTGGGTGTGGCGGACCGCTATCAGGACATAGCGTTGGCTACCCGTGATATTG
CTGAAGAGCTTGGCGGCGAATGGGCTGACCGCTTCCTCGTGCTTTACGGTATCGCCGCTCCCG

ATTCGCAGCGCATCGCCTTCTATCGCCTTCTTGACGAGTTCTTCTGAGCGGGACTCTGGGGTTC
GAAATGACCGACCAAGCGACGCCCAACCTGCCATCACGAGATTTCGATTCCACCGCCGCCTTC
TATGAAAGGTTGGGCTTCGGAATCGTTTTCCGGGACGCCGGCTGGATGATCCTCCAGCGCGGG
GATCTCATGCTGGAGTTCTTCGCCCACCCCAACTTGTTTATTGCAGCTTATAATGGTTACAAATA
AAGCAATAGCATCACAAATTTCACAAATAAAGCATTTTTTTCACTGCATTCTAGTTGTGGTTTG
TCCAAACTCATCAATGTATCTTATCATGTCTGTATACCGTCGACCTCTAGCTAGAGCTTGGCGTA
ATCATGGTCATAGCTGTTTCCTGTGTGAAATTGTTATCCGCTCACAATTCCACACAACATACGA
GCCGGAAGCATAAAGTGTAAAGCCTGGGGTGCCTAATGAGTGAGCTAACTCACATTAATTGCG
TTGCGCTCACTGCCCGCTTTCCAGTCGGGAAACCTGTCGTGCCAGCTGCATTAATGAATCGGC
CAACGCGCGGGGAGAGGCGGTTTGCGTATTGGGCGCTCTTCCGCTTCCTCGCTCACTGACTCG
CTGCGCTCGGTCGTTCGGCTGCGGCGAGCGGTATCAGCTCACTCAAAGGCGGTAATACGGTTA
TCCACAGAATCAGGGGATAACGCAGGAAAGAACATGTGAGCAAAAGGCCAGCAAAAGGCCA
GGAACCGTAAAAAGGCCGCGTTGCTGGCGTTTTTCCATAGGCTCCGCCCCCCTGACGAGCATC
ACAAAAATCGACGCTCAAGTCAGAGGTGGCGAAACCCGACAGGACTATAAAGATACCAGGC
GTTTCCCCCTGGAAGCTCCCTCGTGCGCTCTCCTGTTCCGACCCTGCCGCTTACCGGATACCTG
TCCGCCTTTCTCCCTTCGGGAAGCGTGGCGCTTTCTCATAGCTCACGCTGTAGGTATCTCAGTT
CGGTGTAGGTCGTTCGCTCCAAGCTGGGCTGTGTGCACGAACCCCCCGTTCAGCCCGACCGC
TGCGCCTTATCCGGTAACTATCGTCTTGAGTCCAACCCGGTAAGACACGACTTATCGCCACTG
GCAGCAGCCACTGGTAACAGGATTAGCAGAGCGAGGTATGTAGGCGGTGCTACAGAGTTCTT
GAAGTGGTGGCCTAACTACGGCTACACTAGAAGAACAGTATTTGGTATCTGCGCTCTGCTGAA
GCCAGTTACCTTCGGAAAAAGAGTTGGTAGCTCTTGATCCGGCAAACAAACCACCGCTGGTA
GCGGTGGTTTTTTTGTTTGCAAGCAGCAGATTACGCGCAGAAAAAAAGGATCTCAAGAAGAT
CCTTTGATCTTTTCTACGGGGTCTGACGCTCAGTGGAACGAAAACTCACGTTAAGGGATTTTG
GTCATGAGATTATCAAAAAGGATCTTCACCTAGATCCTTTTAAATTAAAAATGAAGTTTTAAAT
CAATCTAAAGTATATATGAGTAAACTTGGTCTGACAGTTACCAATGCTTAATCAGTGAGGCACC
TATCTCAGCGATCTGTCTATTTCGTTCATCCATAGTTGCCTGACTCCCCGTCGTGTAGATAACTA
CGATACGGGAGGGCTTACCATCTGGCCCCAGTGCTGCAATGATACCGCGAGACCCACGCTCAC
CGGCTCCAGATTTATCAGCAATAAACCAGCCAGCCGGAAGGGCCGAGCGCAGAAGTGGTCCT
GCAACTTTATCCGCCTCCATCCAGTCTATTAATTGTTGCCGGGAAGCTAGAGTAAGTAGTTCGC
CAGTTAATAGTTTGCGCAACGTTGTTGCCATTGCTACAGGCATCGTGGTGTCACGCTCGTCGTT
TGGTATGGCTTCATTCAGCTCCGGTTCCCAACGATCAAGGCGAGTTACATGATCCCCCATGTTG
TGCAAAAAAGCGGTTAGCTCCTTCGGTCCTCCGATCGTTGTCAGAAGTAAGTTGGCCGCAGT
GTTATCACTCATGGTTATGGCAGCACTGCATAATTCTCTTACTGTCATGCCATCCGTAAGATGCT
TTTCTGTGACTGGTGAGTACTCAACCAAGTCATTCTGAGAATAGTGTATGCGGCGACCGAGTT
GCTCTTGCCCGGCGTCAATACGGGATAATACCGCGCCACATAGCAGAACTTTAAAAGTGCTCA
TCATTGGAAAACGTTCTTCGGGGCGAAAACTCTCAAGGATCTTACCGCTGTTGAGATCCAGTT
CGATGTAACCCACTCGTGCACCCAACTGATCTTCAGCATCTTTTACTTTCACCAGCGTTTCTGG
GTGAGCAAAAACAGGAAGGCAAAATGCCGCAAAAAAGGGAATAAGGGCGACACGGAAATG
TTGAATACTCATACTCTTCCTTTTTCAATATTATTGAAGCATTTATCAGGGTTATTGTCTCATGAG
CGGATACATATTTGAATGTATTTAGAAAAATAAACAAATAGGGGTTCCGCGCACATTTCCCCGA
AAAGTGCCACCTGACGTC

**b. Fusion Plasmid Map**

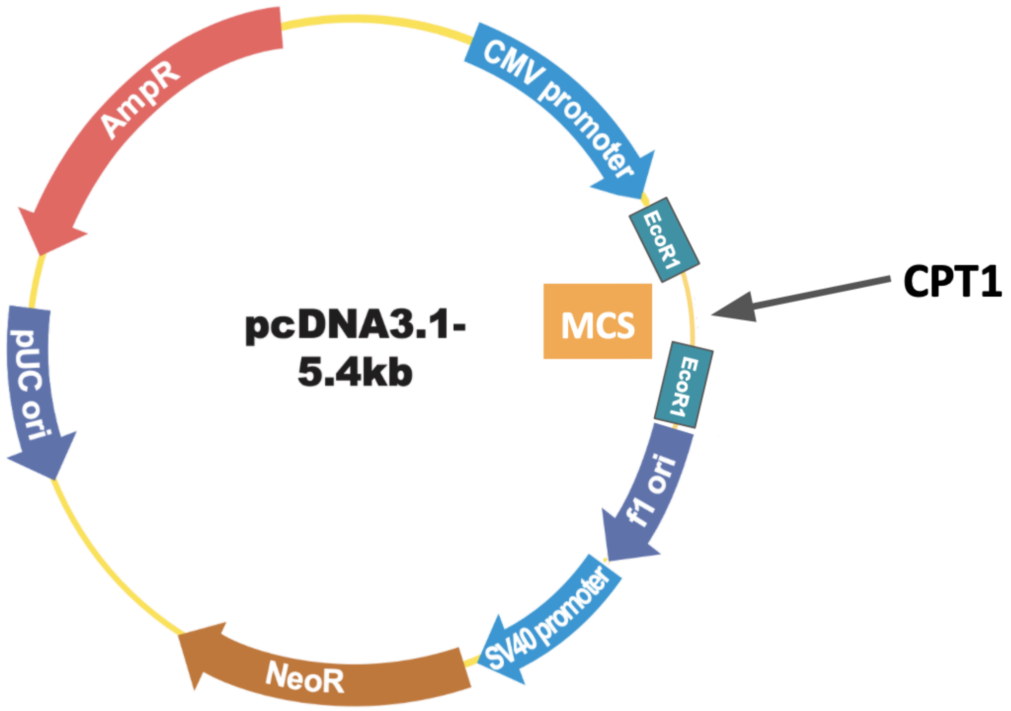

#### C. Standard Curve for DTNB

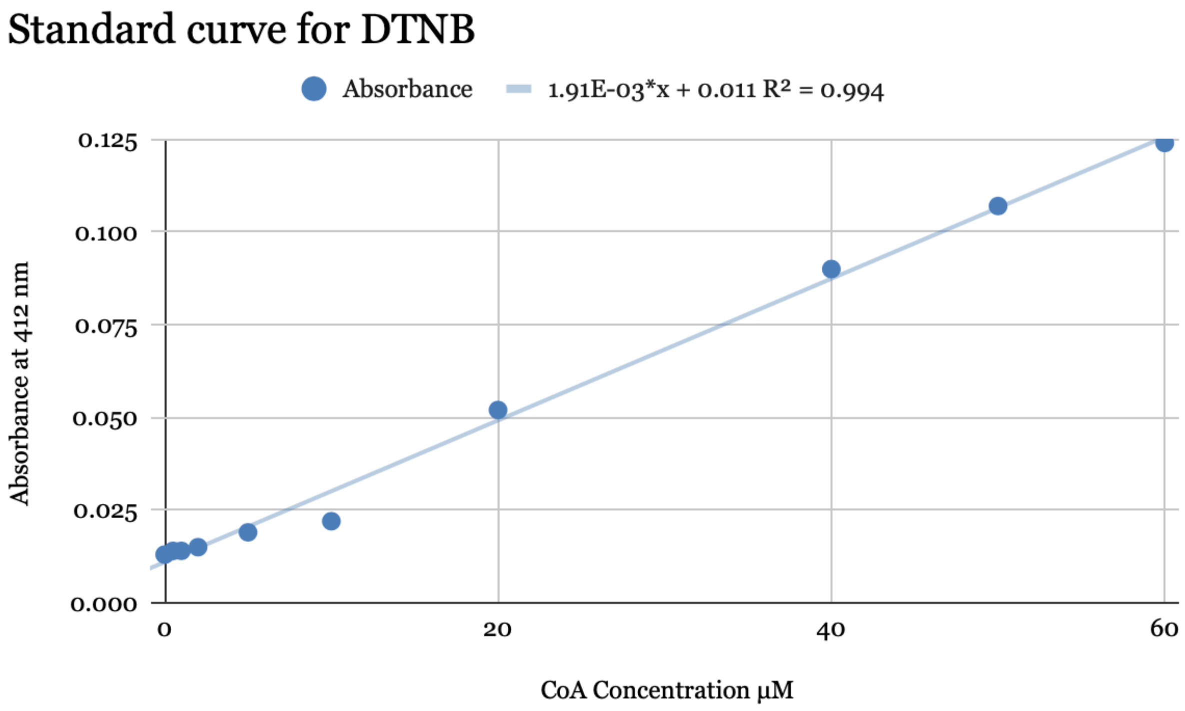

#### D. 78 Compound Library

Compounds employed in the sample high throughput screen include:

| Compound | Effect | Compound | Effect |
|---|---|---|---|
| Isradipine | No inhibition | 5-Fluorouracil | No inhibition |
| Chlorpromazine | Inhibition | Fludrocortisone acetate | No inhibition |
| Paliperidone | No inhibition | Fluocinolone Acetonide | No inhibition |
| Metaxalone | No inhibition | Fludrocortisone acetonide | No inhibition |
| Levodopa | No inhibition | Betamethasone | No inhibition |
| Trichlormethiazide | No inhibition | Hydrocortisone | No inhibition |
| Esomeprazole magnesium | No inhibition | Famotidine | No inhibition |
| Demeclocycline | No inhibition | 3-(2-(tert-butylamino)-1-hydroxyethyl)-2-chlorophenol | No inhibition |
| Carbidopa | No inhibition | Triamcinolone acetonide | No inhibition |
| Vincamine | No inhibition | Itraconazole | No inhibition |
| Berberine chloride | No inhibition | Doxorubicin | No inhibition |
| Pentoxifylline | No inhibition | Donepezil | No inhibition |
| Andrographolide | No inhibition | Pramipexole dihydrochloride | No inhibition |
| Eserine | No inhibition | Oleanolic acid | No inhibition |
| Erythromycin | No inhibition | Palbociclib | No inhibition |
| Mesalamine | No inhibition | Pomalidomide | No inhibition |
| Desloratadine | No inhibition | WZ4002 | No inhibition |

| | | | |
|---|---|---|---|
| Nisoldipine | No inhibition | Paroxetine | No inhibition |
| Felodipine | No inhibition | Motexafin gadolinium | No inhibition |
| Diltiazem HCl | No inhibition | Raloxifene HCl | No inhibition |
| Verapamil HCl | No inhibition | Rapamycin | No inhibition |
| Tolmetin | No inhibition | Camptothecin | No inhibition |
| Nimodipine | No inhibition | SN-38 | No inhibition |
| Carprofen | No inhibition | 5-hydroxypropafenone HCl | No inhibition |
| Taxol | No inhibition | Tulobuterol | No inhibition |
| Podophyllotoxin | No inhibition | Galantamine | No inhibition |
| Papaverine | No inhibition | Ranolazine | No inhibition |
| Ropinirole HCl | No inhibition | Bupivacaine | No inhibition |
| Yohimbine HCl | No inhibition | Imipramine | No inhibition |
| Naringenin | No inhibition | Tramadol | No inhibition |
| Quinine HCl | No inhibition | Primidone | No inhibition |
| Quercetin | No inhibition | Carvedilol phosphate | No inhibition |
| Promethazine | No inhibition | Metoprolol succinate | No inhibition |
| Nifedipine | No inhibition | Pindolol | No inhibition |
| Lacosamide | No inhibition | Metformin | No inhibition |
| Paroxetine HCl | No inhibition | Thalidomide | No inhibition |
| Venlafaxine HCl | No inhibition | Ketoprofen | No inhibition |
| S-(-)carvedilol | No inhibition | Mesalamine | No inhibition |
| Clozapine | No inhibition | | |
